# Recurrent evolution of an *APO1* cis-regulatory module that shapes rice panicle branching

**DOI:** 10.64898/2026.11.18.767244

**Authors:** Takashi Makino, Katsutoshi Tsuda, Asa Makino, Kei Saito, Taiyo Toriba, Junko Kyozuka

## Abstract

Quantitative variation in inflorescence branching drives grass diversification, yet the cis-regulatory mechanisms tuning reproductive meristem activity remain poorly understood. Here, we identified a module of two CArG boxes within a conserved noncoding sequence near rice *APO1*. Genome-edited alleles generated a quantitative phenotypic series: deletions affecting both CArG boxes produced the strongest increases in secondary branching, whereas deletions restricted to CArG box 1 produced smaller but reproducible increases. These deletions extended secondary branch formation toward the panicle base at the expense of direct spikelets, while spikelet number per primary branch was maintained or increased. Across eight genotypes, *APO1* transcript abundance was quantitatively associated with branching, and both CArG boxes contributed to MADS15–MADS34 binding to CNS3, consistent with this module limiting *APO1* expression. Structural and CArG-box changes in this region were detected across multiple phylogenetically distinct lineages with branched inflorescences, indicating that *APO1* underwent recurrent cis-regulatory modification during grass inflorescence diversification.

---

Grass inflorescences range from compact spikes to highly branched panicles. Differences in the initiation, maintenance, and fate transitions of reproductive meristems generate this architectural diversity and determine the number and spatial arrangement of spikelets ^1–3^. Genetic studies have identified conserved regulators of these developmental processes, but much less is known about how evolutionary changes in their cis-regulatory sequences quantitatively alter inflorescence architecture. Because cis-regulatory variation can modify the level, timing, or spatial domain of gene expression without changing gene function, it provides a plausible mechanism for generating graded evolutionary changes in branching ^4^.

*ABERRANT PANICLE ORGANIZATION 1* (*APO1*), which encodes an F-box protein orthologous to *Arabidopsis UNUSUAL FLORAL ORGANS* (*UFO*), is a dosage-sensitive regulator of rice inflorescence development ^5–7^. In rice, loss of *APO1* function causes premature acquisition of spikelet meristem identity and reduces spikelet number, whereas increased *APO1* expression prolongs the inflorescence meristem and branch meristem phases, thereby increasing branching and spikelet production ^5,8,9^. Natural variation at the *APO1* locus also affects panicle architecture and spikelet production ^6,9^. More recently, targeted deletion of a chromatin accessibility-defined region in the *APO1* promoter increased grain yield under field conditions ^10^. Similarly, natural and experimentally generated variation in the wheat ortholog *WAPO1* affects spikelet number per spike ^11^. Yet the specific cis-regulatory motifs that control *APO1* expression, the transcription factors that recognize them, and the extent to which their divergence contributed to grass inflorescence evolution have remained unresolved.

Conserved noncoding sequences (CNSs) are promising candidates for developmental cis-regulatory elements because evolutionary constraint can reveal functional noncoding regions that are difficult to recognize from sequence features in a single species ^12,13^. Comparative genomic studies have identified CNSs associated with gene regulation and agronomic traits in maize and other grasses ^14,15^, while a recent analysis of 284 plant species revealed extensive CNS conservation across deep evolutionary timescales and demonstrated the developmental importance of ancient CNSs ^16^. Integrating sequence conservation, chromatin accessibility, transcription factor binding motifs, and other genomic features has also enabled targeted promoter editing to generate quantitative trait variation in rice ^17^. Editing conserved regulatory regions of meristem regulators can generate allelic series with graded phenotypic effects: targeted editing of a conserved promoter region of the tomato *UFO* ortholog *ANANTHA* revealed antagonistic cis-regulatory activities that stabilize floral development ^18^, and editing a downstream CNS of rice *TAWAWA1* (*TAW1*) generated graded variation in panicle branching ^19^.

MADS-domain proteins are central regulators of inflorescence and floral meristem identity and recognize cis-regulatory sequences known as CArG boxes ^20^. In rice, *MADS34* (*PAP2*) regulates panicle branching and spikelet development ^21,22^, while *MADS14, MADS15, MADS18,* and *MADS34* cooperatively regulate inflorescence meristem identity, and *MADS15* and *MADS34* physically interact ^23^. Whether any of these factors act through defined cis-regulatory elements at the *APO1* locus, however, has not been established. Here, we identify three CNSs surrounding rice *APO1* and characterize two conserved CArG-box motifs within CNS3 that together constitute a cis-regulatory module.

## Results

### Comparative genomics identifies three conserved noncoding sequences near *APO1*

To investigate the cis-regulatory basis of *APO1* expression, we compared the genomic region surrounding the rice *APO1* locus across representative species of Poaceae to identify conserved noncoding sequences. We identified three noncoding sequences, designated CNS1, CNS2, and CNS3, that were conserved between rice and other grasses (Fig. 1a–c and Supplementary Fig. 1). CNS1 showed broad sequence conservation across the sampled Poaceae, whereas CNS2 was less strongly conserved. CNS3 contained discrete, highly conserved segments shared across the BEP and PACMAD clades. No homologous sequences meeting our detection criteria were identified in the syntenic region of *Musa acuminata*, suggesting that detectable conservation of these three elements is restricted to grasses at the phylogenetic resolution of our analysis.

**Fig. 1.**
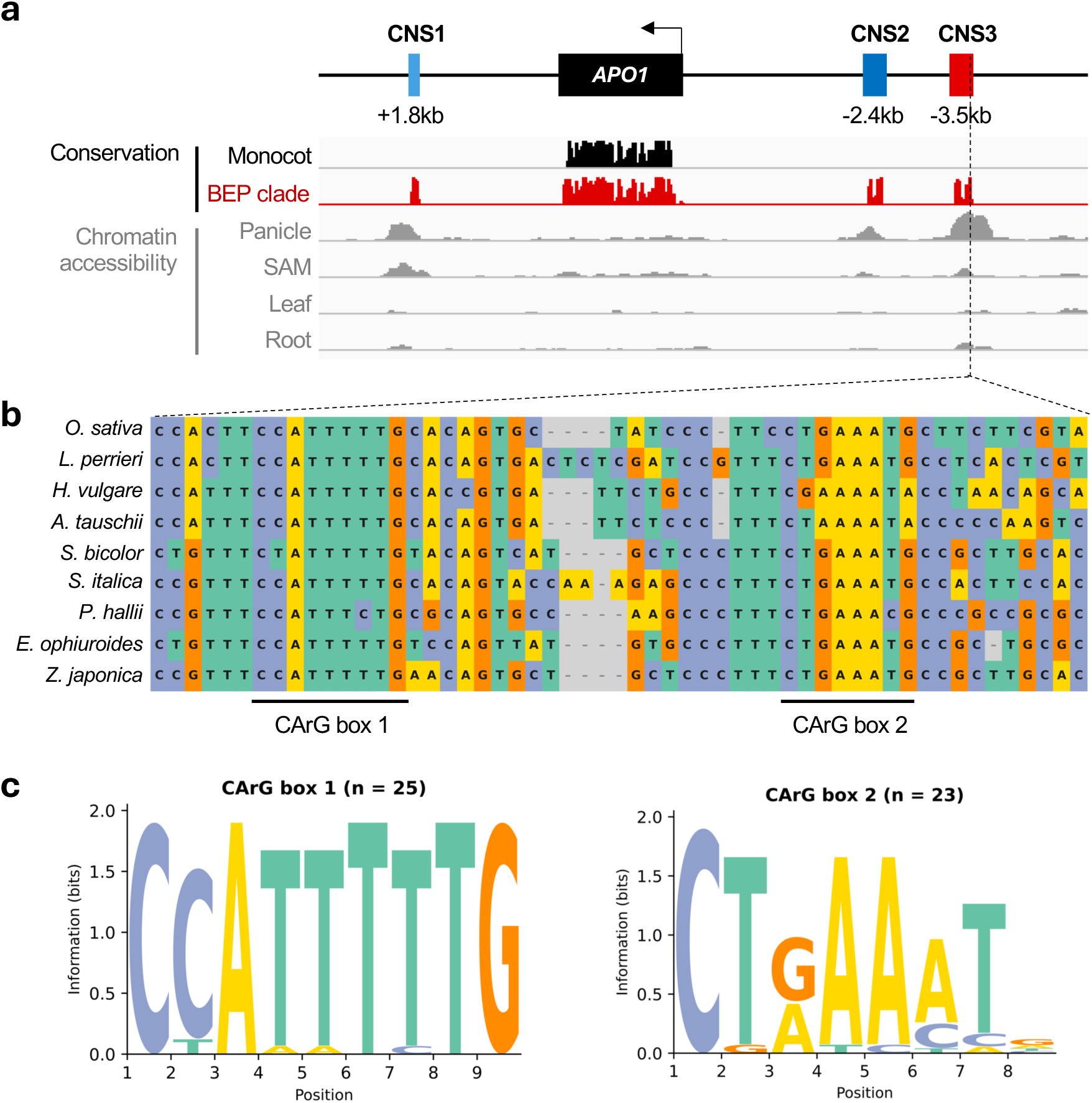
A cis-regulatory module with paired CArG boxes flanks rice *APO1*. **a,** Genomic organization of the rice *APO1* locus and chromatin accessibility across the regulatory region. Upper: the *APO1* gene model and transcriptional orientation, CNS1, CNS2, and CNS3 with their distances from *APO1*, and Poaceae conservation tracks (Monocot and BEP clade). Lower: chromatin-accessibility profiles from the CART dataset ^24^ in panicle, SAM, leaf, and root tissues. **b,** Multiple-sequence alignment of the CArG box 1 and CArG box 2 intervals within CNS3 across representative Poaceae species. Conserved positions, substitutions, indels, and motif boundaries are indicated. Full-length CNS alignments for representative species are shown in Supplementary Fig. 1. **c,** Sequence logos of CArG box 1 (n = 25) and CArG box 2 (n = 23) across the analyzed Poaceae species. Height at each position represents information content in bits. Species for which a motif was absent or undetectable were excluded.

CNS1, CNS2 and CNS3 overlapped accessible chromatin regions in developing rice panicles in the Chromatin Accessibility of Rice Tissues (CART) dataset. CNS1 also showed accessibility in Shoot Apical Meristem (SAM), whereas CNS2 and CNS3 showed low or undetectable accessibility in the other tissues examined (Fig. 1a) ^24^. This panicle-associated accessibility identifies the CNSs as candidate cis-regulatory elements active during inflorescence development.

Sequence inspection of CNS3 identified two closely spaced CArG-box motifs recognized by MADS-domain transcription factors, designated CArG boxes 1 and 2 (Fig. 1b). Sequence-logo analysis revealed that CArG box 1 was nearly invariant across the sampled Poaceae, whereas CArG box 2 was more variable (Fig. 1c).

### Editing the *APO1* regulatory region produces a quantitative series of branching phenotypes

To assess the functions of the three CNSs surrounding *APO1*, we generated 34 independent homozygous lines carrying CRISPR-induced mutations within or around CNS1, CNS2, and CNS3 (Fig. 2a and Supplementary Fig. 2; Supplementary Table 1). We quantified primary branch number, secondary branch number, total spikelet number, and lateral-organ number (secondary branches + spikelets directly borne on primary branches; Fig. 2b). To account for differences in overall panicle size, we also evaluated secondary branches per lateral organ as a measure of relative secondary branching.

**Fig. 2.**
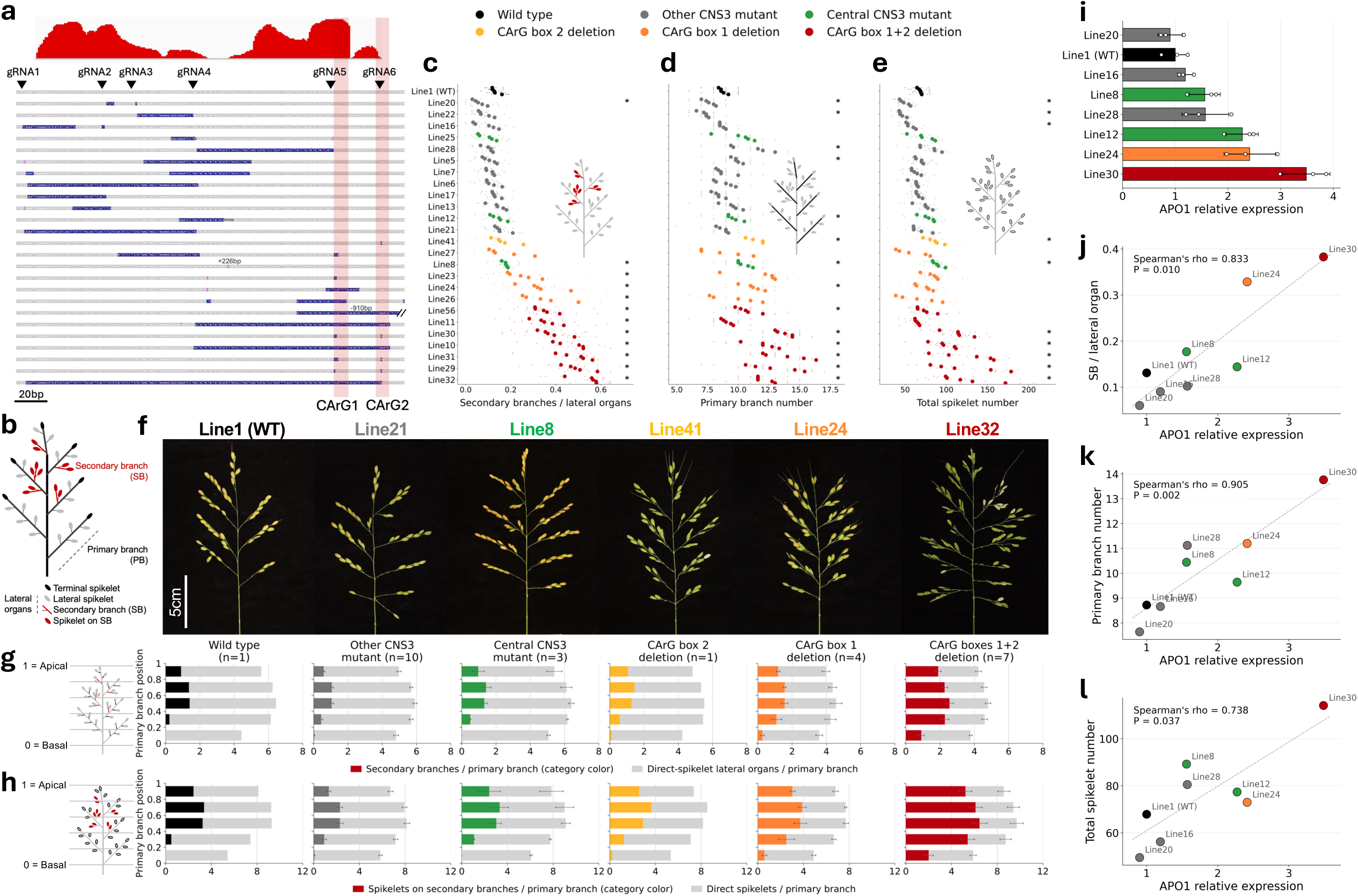
CNS3 editing alters panicle branching, lateral organ fate, and APO1 expression. **a,** CNS3 edited alleles showing phastCons conservation scores, CArG boxes 1 and 2, and guide RNA target sites (inverted triangles). Each bar represents a homozygous line; deletions (dark blue), substitutions (sequence colors), insertions (pink), and inversions (purple) are indicated. **b,** Panicle architecture showing primary branches (PBs), secondary branches (SBs), and terminal (black), lateral (gray), and SB-borne (red) spikelets. **c,** secondary branches per lateral organ, **d,** primary branch number, and **e,** total spikelet number. Small open circles indicate individual panicles and large filled circles plant means (up to five panicles from each of five plants per line). Plant means were used as independent biological replicates. For **c,** ratios were calculated for each primary branch and averaged within panicles. Plants within each line are arranged in ascending trait order; lines are ordered by mean secondary branches per lateral organ and colored by mutation class. Asterisks indicate significant differences from wild type (Mann–Whitney U test, Benjamini–Hochberg-adjusted p < 0.05; exact values in Supplementary Table 2). Phenotypes of CNS1, CNS2, and CNS2–CNS3 intergenic mutants are shown in Supplementary Fig. 2. **f,** Representative panicles from lines 1 (wild type), 21 (other CNS3 mutant), 8 (central CNS3 mutant), 41 (CArG box 2 deletion), 24 (CArG box 1 deletion), and 32 (CArG box 1+2 deletion). Scale bar, 5 cm. g, h, Spatial distributions of lateral organs (g) and spikelets (h) across five normalized primary-branch positional bins (0, basal; 1, apical). Bars distinguish secondary branches (colored) from direct-spikelet lateral organs (gray) in **g,** and SB-borne (colored) from direct (gray) spikelets in **h.** Terminal spikelets were excluded from g but included in h. Mutation-class values are unweighted means of line means, with error bars indicating SEM among line means for components and totals. Colors indicate mutation classes, and numbers of independent lines are shown. i, *APO1* expression measured by qRT-PCR and normalized to OsUBQ5 (n = 3 biological replicates per genotype). Points indicate replicates; bars show mean ± SD. **j,** Genotype-mean *APO1* expression versus secondary branches per lateral organ; **k,** primary branch number; and **l,** total spikelet number. Points represent genotypes (n = 8), labeled by line number and colored by mutation class. Dashed lines guide the eye; Spearman’s ρ and P values are indicated.

Mutations in CNS1 and CNS2 consistently reduced secondary branches per lateral organ and total spikelet number, although these reductions were statistically significant in only some lines (Supplementary Fig. 2 and Supplementary Table 2). In contrast, CNS3 mutations generated a clear allelic series (Fig. 2c–f). Deletions encompassing both CArG boxes significantly increased secondary branches per lateral organ in all seven lines (10, 11, 29, 30, 31, 32, and 56), with a 3.4-fold increase on average relative to wild type (Fig. 2c; Supplementary Table 2). Primary branch number and total spikelet number increased by 1.39-fold and 1.52-fold on average, respectively, with significant increases in six and five of the seven lines (Fig. 2d, e). Line 11 showed comparatively weak effects on these two traits, with neither reaching statistical significance.

Deletions restricted to CArG box 1 (lines 23, 24, 26, and 27) produced smaller increases in relative secondary branching than deletions affecting both CArG boxes. These increases were statistically significant in lines 23, 24, and 26 but not in line 27 (Fig. 2c; Supplementary Table 2). Primary branch number increased significantly only in line 24, and total spikelet number did not increase significantly in any of these lines. Thus, CArG box 1 deletions predominantly affected relative secondary branching rather than overall panicle expansion.

The sole CArG box 2 deletion line (41) showed a distinct phenotype: primary branch number and total spikelet number increased significantly, whereas relative secondary branching did not (Fig. 2c–e; Supplementary Table 2).

Additional CNS3 alleles altered the central region while retaining both CArG boxes: a 226-bp insertion in line 8, a 28-bp deletion with a 6-bp substitution in line 12, and a complex local rearrangement in line 25 (Fig. 2a; Supplementary Table 1). Line 8 showed significant increases in primary branch number, total spikelet number, and relative secondary branching, resembling the dominant *apo1-D* phenotype (Fig. 2c–e) ^9^. Line 12 significantly increased primary branch number alone, whereas line 25 showed no significant increase in the branching traits examined. Larger central-region deletions retaining both CArG boxes (lines 5, 6, and 7) did not produce comparable increases: relative secondary branching and total spikelet number showed no significant differences from wild type, and only line 5 showed a significant increase in primary branch number. These contrasting effects indicate that CNS3 regulatory output depends on local structural context as well as CArG-box integrity.

Deletions between CNS2 and CNS3 also produced distinct phenotypic effects. Smaller deletions of 143–160 bp (lines 34 and 37) did not significantly affect relative secondary branching or total spikelet number, although primary branch number increased in line 37 (Supplementary Fig. 2). In contrast, a 703-bp deletion (line 36) significantly reduced relative secondary branching while increasing primary branch number (Supplementary Table 2).

No tertiary branches were observed in any of the 821 panicles from 175 plants across 35 lines examined. Thus, even the strongest CNS3 mutations increased branching only at the secondary-branch level.

### CNS3 editing shifts the spatial domain and fate of lateral organ formation

We next examined the spatial distribution of secondary branches and the allocation of lateral organs between direct spikelets and secondary branches. Primary branch positions were normalized from 1 (most apical) to 0 (most basal) to compare panicles with different numbers of primary branches.

In wild type panicles, secondary branches were concentrated in mid-apical positions, with none on the most basal primary branch (Fig. 2g). Deletions encompassing both CArG boxes expanded secondary branch formation toward the panicle base, including the most basal primary branches. The mean onset position of secondary branching shifted from 0.47 in wild type to 0.17 in both-box deletion lines, with significant shifts in all seven independent lines (Supplementary Table 3). CArG box 1 deletions produced an intermediate shift (mean 0.35), significant only in line 24.

Figure 2g and Supplementary Fig. 3 also show differences in the total number of lateral organs per primary branch among genotypes. In wild type plants, the number of lateral organs per primary branch was similar across most panicle positions and lower on the most basal primary branches, a pattern generally observed across genotypes. In lines with deletions affecting both CArG boxes, the total number of lateral organs per primary branch was reduced across all positions, while the secondary-branch component expanded as the direct-spikelet component contracted. This pattern indicates that the increase in secondary branch number does not reflect production of additional lateral organs, but rather a shift in lateral-organ fate from direct spikelets toward secondary branches. Deletions restricted to CArG box 1 showed an intermediate reduction in total lateral organ number and an intermediate expansion of secondary branch fate, between wild type and the phenotypes of lines with deletions affecting both CArG boxes.

We next examined the spatial distribution of spikelet production across all mutation classes (Fig. 2h and Supplementary Fig. 3). Although lines with deletions affecting both CArG boxes reduced the number of lateral organs per primary branch, total spikelet production was maintained or increased (Fig. 2h): At the most basal primary branch, total spikelet numbers were similar to wild type (5.9 vs. 5.4), but approximately two spikelets were borne on secondary branches in the deletion lines (Fig. 2h). At mid-panicle positions, secondary-branch-borne spikelets increased (6.5 vs. 3.2), compensating for fewer direct spikelets (3.2 vs. 6.0) and maintaining slightly higher total spikelet numbers (9.7 vs. 9.3).

Central CNS3 mutants (lines 8, 12, and 25) showed a distinct phenotypic profile. These lines tended to increase primary branch number and, in line 8, total spikelet number (Fig. 2d, e), but showed little or no shift in secondary branch onset position (Supplementary Table 3). Line 8 showed a modest but statistically significant increase in secondary branches per lateral organ, whereas lines 12 and 25 did not (Supplementary Table 2).

### *APO1* transcript abundance is associated with the degree of secondary branching

To examine the association between *APO1* expression and branching phenotypes, we quantified *APO1* expression in 1–2-mm young panicles from eight genotypes spanning the observed phenotypic range by RT-qPCR (Fig. 2i). Expression was lowest in line 20 and highest in line 30. Genotype-level mean *APO*1 expression was positively correlated with primary branch number (Spearman’s rho = 0.905, P = 0.002; Fig. 2k), secondary branches per lateral organ (rho = 0.833, P = 0.010; Fig. 2j), and total spikelet number (rho = 0.738, P = 0.037; Fig. 2l; n = 8 genotypes, three biological replicates per genotype).

### Both CArG box 1 and CArG box 2 contribute to MADS15–MADS34 binding to CNS3

CNS3 contains two conserved CArG-box motifs that may be recognized by MADS-domain factors. *APO1* expression declines during early inflorescence development as several MADS-box genes, including *MADS14*, *MADS15*, *MADS18*, and *MADS34*, increase in expression ^23,25^. Previous studies have shown that MADS34 regulates rice inflorescence branching ^21^ and directly binds CArG-box-containing regulatory regions of RCN4 ^22^. Notably, MADS15 and MADS34 have been shown to physically interact ^23^. We therefore tested their binding to the CNS3 region containing CArG boxes 1 and 2 by electrophoretic mobility shift assays (EMSA; Fig. 3). MADS14 and MADS15 produced weak shifts individually, whereas MADS18 and MADS34 showed no detectable binding alone. Among the tested combinations, MADS14–MADS34 and MADS15–MADS34 formed detectable complexes, with MADS15–MADS34 showing the strongest binding (Fig. 3b).

**Fig. 3.**
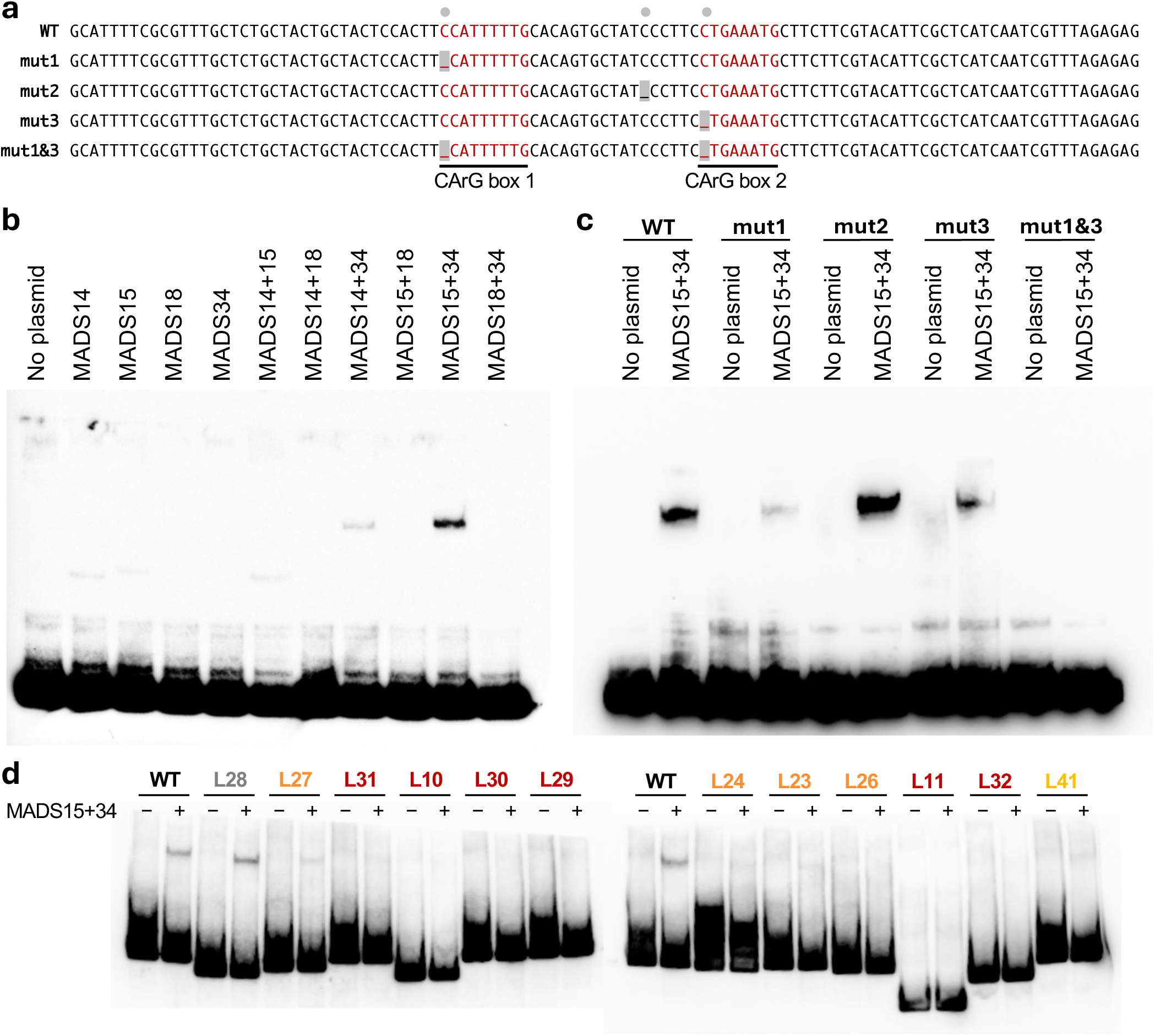
MADS15–MADS34 binding to *APO1* CNS3 is sensitive to disruption of the two CArG boxes. **a,** CNS3 probe design. Wild type and mutant probes used to examine the contribution of the two CArG boxes to MADS15–MADS34 binding are shown. The mutant probes contain a single-nucleotide deletion in CArG box 1 (mut1), a single-nucleotide deletion in the sequence between the two CArG boxes (mut2), a single-nucleotide deletion in CArG box 2 (mut3), or the combined mut1 and mut3 deletions (mut1&3). CArG-box sequences are shown in red, deleted nucleotides are indicated in gray, and gray circles above the sequences mark the positions of the introduced deletions. **b,** EMSA of MADS14, MADS15, MADS18, and MADS34 with the wild type CNS3 probe. The proteins were assayed individually and in the indicated combinations. MADS15–MADS34 produced the strongest shifted signal among the combinations tested. **c,** EMSA of the wild type, mut1, mut2, mut3, and mut1&3 CNS3 probes with MADS15 and MADS34. Single-nucleotide deletion of either CArG box reduced but did not abolish detectable binding, whereas mut2 retained binding. Simultaneous deletion of both CArG boxes (mut1&3) abolished detectable binding under the assay conditions. **d,** EMSA using CNS3 probes corresponding to the actual genome-edited alleles. Probes derived from the wild type and edited lines 28, 27, 31, 10, 30, 29, 24, 23, 26, 11, 32, and 41 were assayed in the presence or absence of MADS15 and MADS34. The wild type and line 28 probes showed clear shifted signals, whereas line 27 showed weak residual binding. Detectable binding was abolished in lines 23, 24, and 26 carrying CArG-box-1 deletions, line 41 carrying a CArG-box-2 deletion, and lines 10, 11, 29, 30, 31, and 32 carrying deletions affecting both CArG boxes.

Single-nucleotide deletion of CArG box 1 (mut1) or CArG box 2 (mut3) reduced but did not abolish MADS15–MADS34 binding, whereas simultaneous deletion of both CArG boxes (mut1&3) abolished detectable binding (Fig. 3a, c). A single-nucleotide deletion between the motifs (mut2) did not reduce binding.

We next tested CNS3 fragments amplified from the edited lines used for phenotypic analysis (Fig. 3d). Wild type CNS3 and line 28, which retained both intact CArG boxes, showed detectable binding. Binding was abolished in lines with deletions affecting both CArG boxes (lines 10, 11, 29, 30, 31, and 32), CArG box 2 (line 41), or CArG box 1 (lines 23, 24, and 26). Line 27, carrying a CArG box 1 deletion, retained weak binding, consistent with its mild phenotype. These results demonstrate that CArG-box-dependent binding occurs not only with synthetic probes but also with CNS3 fragments derived from the edited lines.

Together, these biochemical and genetic results support a model in which MADS15–MADS34 binding to CNS3 limits *APO1* expression and restricts secondary branch formation to mid-apical primary branches (Fig. 5).

### The *APO1* regulatory module changed recurrently during grass inflorescence evolution

Grass inflorescences vary widely in architecture, ranging from unbranched spikes and racemes to highly branched panicles (Fig. 4a). To determine whether the *APO1* regulatory region shows phylogenetically correlated variation with this diversity, we surveyed the CNS3 interval and its flanking sequences across grass species with chromosome-scale genome assemblies and mapped regulatory structure and inflorescence architecture onto the species phylogeny (Fig. 4b and Supplementary Table 4).

**Fig. 4.**
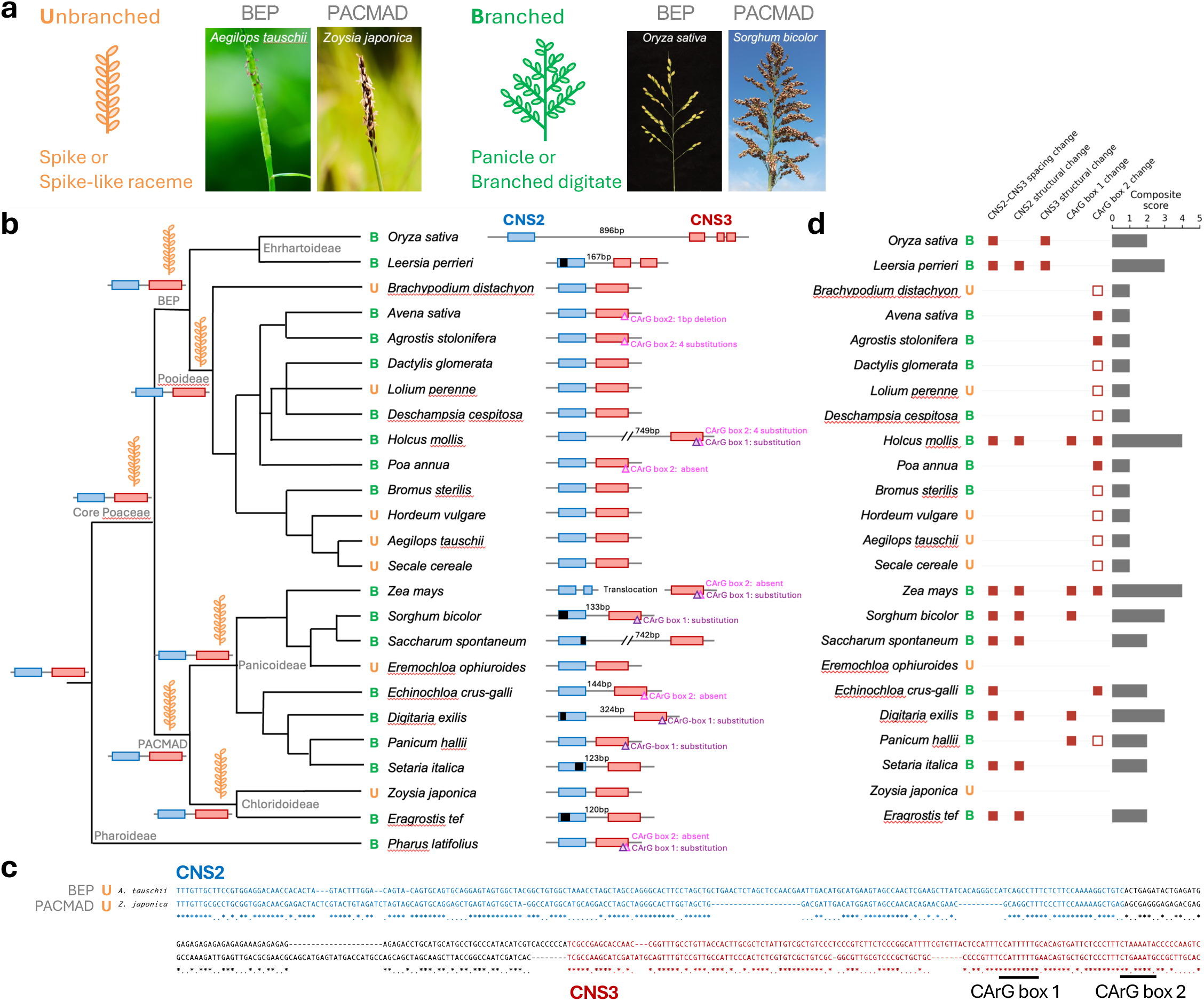
Recurrent modification of the *APO1* cis-regulatory module during grass inflorescence evolution. **a,** Diagrams and photographs of unbranched (spike or spike-like raceme) and branched (panicle or branched digitate) inflorescences from the BEP and PACMAD clades. Photographs were cropped to highlight inflorescence architecture. The *Oryza sativa* photograph was taken by the authors; others were obtained from Wikimedia Commons: *Aegilops tauschii*, Vengolis (CC BY-SA 3.0); *Sorghum bicolor*, Stefan.lefnaer (CC BY-SA 4.0); and *Zoysia japonica*, bastus917 (CC BY-SA 2.0). **b,** Species phylogeny showing inflorescence architecture and *APO1* cis-regulatory structures. Internal branch symbols and CNS structures represent putative ancestral states based on parsimony. Orange and green indicate unbranched and branched architectures, respectively. Extant species are labeled U (unbranched) or B (branched). CNS2 (blue) and CNS3 (red) are positioned according to their intergenic distances. Black segments indicate internal CNS deletions or terminal regions lacking detectable similarity (potential deletion or sequence divergence). Purple and pink triangles indicate changes in CArG boxes 1 and 2, respectively. Structural changes, translocations, and intergenic distances are annotated. **c,** Alignment of the *APO1* cis-regulatory region between the unbranched BEP species *Aegilops tauschii* and PACMAD species *Zoysia japonica*. CNS2 (blue), CNS3 (red), and both CArG boxes are highlighted; asterisks indicate identical nucleotides. **d,** Changes in five regulatory features relative to the inferred ancestral state: CNS2–CNS3 spacing, CNS2 structure, CNS3 structure, CArG box 1, and CArG box 2. Red squares indicate changes; absence of a square indicates conservation. For CArG box 2, filled squares represent major changes (≥4 substitutions, deletion, or motif absence) and open squares minor changes (1–3 substitutions). Gray bars show the composite divergence score (number of changed features, 0–5). U and B indicate unbranched and branched species.

**Fig. 5.**
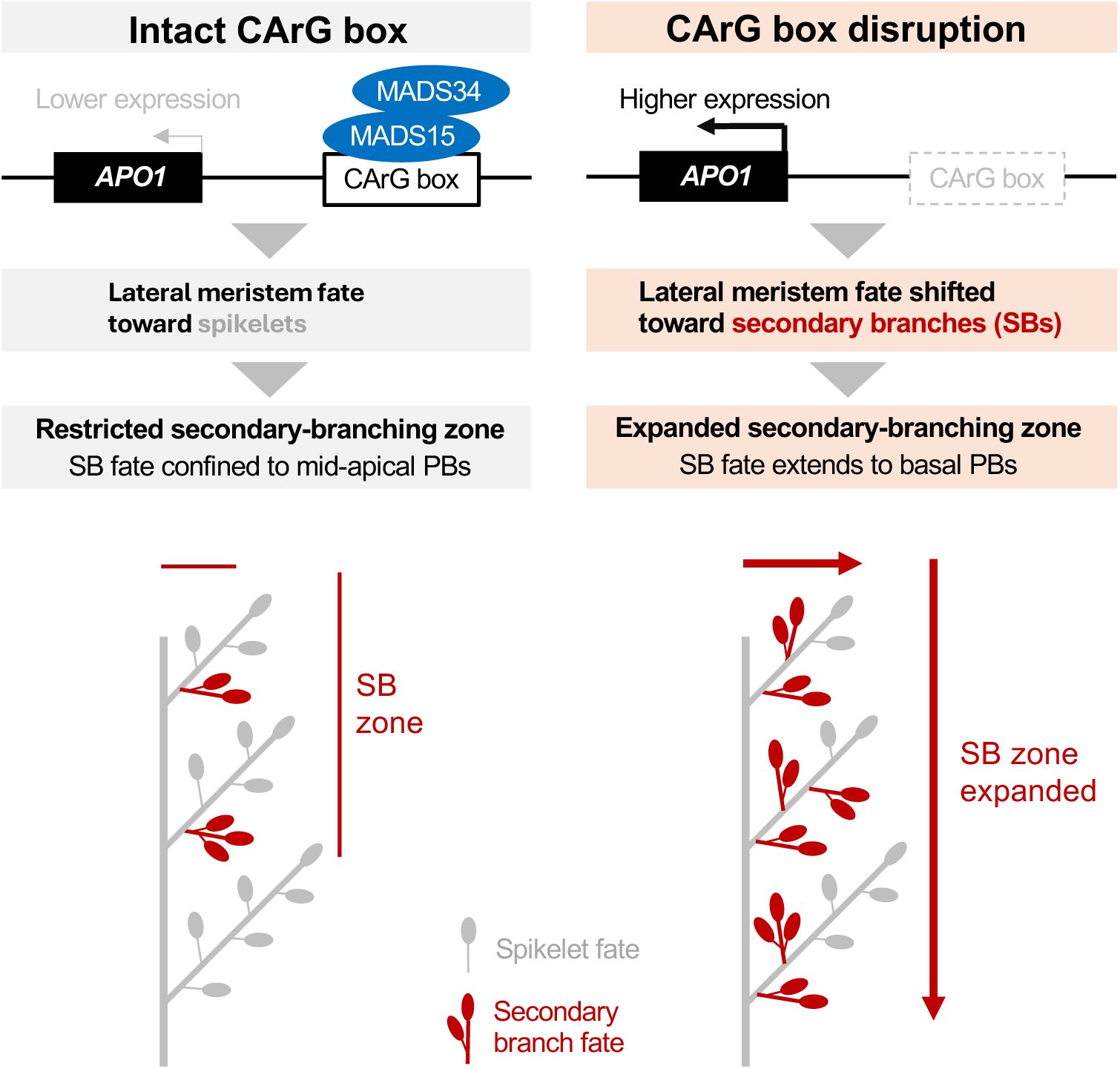
Model linking the CNS3 CArG-box module to the spatial domain of secondary-branch formation. MADS15–MADS34 binding to intact CArG boxes is associated with lower *APO1* expression and restriction of secondary-branch fate to mid-apical primary branches. CArG-box disruption reduces MADS15–MADS34 binding and is associated with higher *APO1* expression, shifting lateral meristem fate toward secondary branches and expanding the secondary-branching zone toward basal primary branches.

CArG box 1 was nearly invariant across the sampled Poaceae, with substitutions detected only in a few species. In contrast, CArG box 2 was more variable, with substitutions in both spike-and panicle-bearing lineages (Fig. 1c). Thus, substitutions in CArG box 2 alone were not consistently associated with increased branching. Changes in CArG box 1 or loss of CArG box 2 were concentrated in more highly branched lineages, whereas many spike-bearing taxa retained conserved motifs (Fig. 4b, Supplementary Fig. 4a and Supplementary Table 5).

Structural changes were also observed in multiple lineages, including CNS2 deletions, CNS3 insertions, insertions between CNS2 and CNS3, and translocation of CNS2 to a different chromosome. These changes occurred predominantly in panicle-bearing lineages, whereas spike-bearing taxa generally retained an intact CNS configuration (Fig. 4b, c, Supplementary Table 6). To quantify these associations, we scored five cis-regulatory features across 24 species: CNS2–CNS3 spacing change (>110 bp or translocation), CNS2 structural change (>30 bp), CNS3 structural change (>30 bp), CArG box 1 substitution, and CArG box 2 change (nucleotide substitution relative to the ancestral CTGAAATG motif, deletion, or motif absence). Branched species showed higher composite divergence scores even after accounting for phylogenetic non-independence (phylogenetic GLS, β = 1.42, SE = 0.46, p = 0.0050; Fig. 4d and Supplementary Table 7). Because minor substitutions in CArG box 2 were observed across many grass species, we also tested a composite score that excluded these minor changes from the CArG box 2 component. This association remained significant when the composite divergence score was calculated using only major CArG box 2 changes (≥4 nucleotide substitutions, deletion, or motif absence; β = 1.64, SE = 0.50, p = 0.0036; Supplementary Table 7). Among individual features, CNS2–CNS3 spacing change (phylogenetic GLS, p = 0.0060) and CNS2 structural change (p = 0.018) were significantly associated with inflorescence type (Supplementary Table 7).

CArG box 1 was conserved across spike-bearing taxa except in the *Secale cereale* reference assembly, which contained an insertion within the motif. Sanger sequencing of an independent *S. cereale* accession recovered an intact motif (Supplementary Fig. 4b). Because the insertion was not detected by Sanger sequencing, we treated CArG box 1 as intact in *S. cereale* for the comparative analysis.

The CNS2–CNS3 configuration in distantly related unbranched lineages, including *Aegilops tauschii* and *Zoysia japonica*, suggested ancestral conservation (Fig. 4c). Under a parsimonious interpretation, these observations supported an intact, closely linked CNS2–CNS3 configuration as the ancestral state of core Poaceae, followed by independent changes in multiple panicle-bearing lineages (Fig. 4b). Although CNS2 and CNS3 were undetectable in the earliest-diverging sampled grass, *Streptochaeta angustifolia*, *Pharus latifolius*, the earliest-diverging lineage with detectable CNSs, retained a compact 65 bp CNS2–CNS3 interval. This supports an ancestral closely linked configuration extending beyond core Poaceae. The association was not absolute: some panicle-bearing taxa retained the conserved CNS configuration. *APO1* regulatory changes were recurrently associated with increased branching but did not uniquely determine inflorescence type.

## Discussion

The graded CNS3 allelic series links a two-CArG-box module to quantitative changes in *APO1* expression and secondary branching. Deletions affecting both motifs consistently produced the strongest increase in secondary branching, whereas CArG box 1 deletions produced smaller, reproducible effects. The sole CArG box 2 deletion allele (line 41) instead increased primary branch number without a significant increase in relative secondary branching. This distinction is intriguing but requires additional CArG box 2 alleles for confirmation. Importantly, *APO1* promotes both inflorescence meristem growth and the maintenance of branch meristem identity ^8,9^. Thus, the contrasting single-box phenotypes suggest a possible cis-regulatory basis for separating these established *APO1* functions across developmental stages, although this interpretation remains to be tested directly.

EMSA using probes from the edited alleles reinforced the relationship between CArG-box integrity and binding. Line 28, which retained both motifs, bound MADS15–MADS34 comparably to wild type; binding was abolished in the tested alleles disrupting both motifs or either motif alone, except for weak residual binding in line 27. In contrast to the synthetic single-nucleotide deletion probes, most edited alleles removed multiple bases, plausibly accounting for the stronger loss of binding. Line 27 carries a three-nucleotide deletion but retains the sequence CTTTTTTTG, differing at only one position from the CC(A/T)6G consensus; this may explain both residual binding and its mild phenotype. The preferential binding of MADS15–MADS34 to CNS3 is consistent with evidence that the intervening domain of plant MADS transcription factors contributes to dimerization partner selection and DNA-binding specificity ^26^. *MADS5* and *MADS34* have been shown to limit rice panicle branching by directly repressing *RCN4* ^22^. Our findings suggest that MADS15–MADS34 may similarly restrict secondary branching through the *APO1* CNS3 module, although direct transcriptional repression of *APO1* remains to be demonstrated.

Earlier transgenic experiments, conducted before CNS3 was identified, provide retrospective support for this model. A genomic *APO1* construct containing the region now recognized as CNS3 (*pro5kb-APO1*) rescued the *apo1-3* phenotype, whereas a shorter construct lacking CNS3 (*pro3kb-APO1*) increased *APO1* expression and branching, albeit with abnormalities in distal panicle development ^8^. Viewed in light of our findings, these results are consistent with a role for CNS3 in constraining *APO1* expression. In addition, a deletion upstream of CNS3, in a region absent from *pro3kb-APO1*, likewise increased *APO1* expression ^10^, suggesting additional regulatory sequences outside CNS3.

Positional analysis shows that CNS3 regulates the basal extent of secondary-branch formation and lateral-organ fate without necessarily reducing spikelet output. The finding that tertiary branches were absent from all *APO1* CNS3 editing lines, despite marked increases in secondary branch number, distinguishes the role of *APO1* from that of *TAWAWA1* (*TAW1*). *TAW1* promotes inflorescence meristem activity and suppresses the phase transition from branch meristems to spikelet meristems; its gain-of-function alleles and CNS-deletion lines produce tertiary branches, reflecting a generalized extension of meristem branching capacity ^19,27^. By contrast, the *APO1* CNS3 phenotypes identified here highlight a different step: the fate of lateral organs on established primary branches, determining whether each lateral organ primordium becomes a spikelet or a secondary branch meristem. *APO1* and *TAW1* thus provide independent regulatory entry points into the inflorescence developmental program, appearing to act at distinct steps in the sequential elaboration of the grass panicle.

The spatial profile also provides a developmental interpretation for the positional specificity of the wild type phenotype. In wild type rice, secondary branches form preferentially on mid-apical primary branches, suggesting positional differences in the competence to form secondary branches. The basal expansion of secondary branch formation in CArG-box deletion lines is consistent with elevated *APO1* expression extending this competence to more basal primary branch meristems before the transition to spikelet identity occurs. This interpretation aligns with the established role of *APO1* in delaying meristem phase transitions ^8,9^ and places the CNS3 module as a cis-regulatory brake on this delay.

Line 8 carries a 226-bp insertion, line 12 a 28-bp deletion with a 6-bp substitution, and line 25 a local replacement involving a duplicated upstream sequence; their branching effects differ despite their proximity. Importantly, this central region overlaps the clustered transposon insertion sites of the four independently isolated dominant *apo1-D* alleles ^9^. The line 8 insertion coincides with *apo1-D2* and lies one nucleotide from *apo1-D1*, whereas the edits in lines 12 and 25 overlap the *apo1-D4/Ur1* insertion site (Supplementary Fig. 5). Thus, two complementary approaches, *apo1-D* mutant isolation and genome editing, independently identify the same relatively weakly conserved segment between strongly conserved CNS3 regions as functionally sensitive.

Line 8 most closely resembled the dominant *apo1-D* phenotype, increasing primary and secondary branching and spikelet number, and also showed elevated *APO1* expression (Fig. 2i). Its insertion may alter regulatory spacing or chromatin organization. Crucially, larger deletions of the central region in lines 5, 6, and 7, which retained both CArG boxes and, in lines 5 and 7, removed all four *apo1-D* insertion sites, did not produce comparable increases in branching. This contrast argues that the line 8 phenotype cannot be explained by simple loss of central-region sequence and instead may depend on the particular structural consequences of the insertion. More broadly, the differing effects of these alleles suggest that CNS3 is not a simple two-element switch: regulatory output depends on local structural context as well as CArG-box integrity. Separating spacing from sequence effects will require targeted follow-up experiments.

CArG box 1 was nearly invariant among spike-bearing taxa, whereas CArG box 2 varied in both spike- and panicle-bearing lineages. This asymmetry in conservation mirrors the asymmetry in phenotypic effects observed in the rice allelic series: deletions restricted to CArG box 1 generally increased secondary branching, whereas the single CArG box 2 deletion showed a distinct, trait-specific phenotype.

The ancestral inflorescence form of Poaceae remains debated. Classical morphological models proposed a panicle-like branching system as the ancestral condition, from which simpler inflorescence architectures arose largely through reductive processes ^28^, a view also adopted in subsequent studies of grass inflorescence evolution ^29^. In contrast, phylogenetic and developmental analyses have emphasized that superficially similar branched inflorescences can arise from distinct developmental and evolutionary trajectories ^30^. Notably, spike-bearing species in both the BEP and PACMAD clades retain an intact, closely linked CNS2–CNS3 configuration (Fig. 4b, c). Under a parsimonious interpretation, these observations support an intact, closely linked CNS2–CNS3 configuration in the ancestor of core Poaceae, consistent with the possibility that its inflorescence was spike-like. Recurrent modification of this regulatory configuration may have provided a route to increased inflorescence branching in descendant lineages (Fig. 4b). The uncertain status of CNS2 and CNS3 in *Streptochaeta*, where the elements may have been lost or diverged beyond recognition, limits inference concerning the deeper ancestor of all Poaceae but does not affect the reconstructed state for the core Poaceae ancestor.

Independent insertions within CNS3 or between CNS2 and CNS3 occur in panicle-bearing *Oryza sativa*, *Leersia perrieri*, and *Saccharum spontaneum*. Their locations recall the dominant *apo1-D* transposon insertions, which increased *APO1* expression and branching ^9^, suggesting that structural disruption may represent one recurrent route to altered regulation. However, some branched Pooideae species retain an intact CNS2–CNS3 configuration, indicating that structural modification of this module is not required for the evolution of branched inflorescences. The effect of a deletion in the intergenic region between CNS2 and CNS3 in rice further suggests that structural changes in this region can alter panicle branching. Together, these findings refine the model: *APO1* regulatory output may be modified through distinct changes in motif sequence, module organization, or spacing, rather than through a single structural route. The comparative associations do not establish causation in other grasses.

Several questions remain unresolved: direct repression and in vivo MADS15–MADS34 occupancy at CNS3 have not been demonstrated; central-region edits do not isolate spacing effects from sequence effects; and natural CNS configurations have not been functionally tested in their native genetic backgrounds.

The allelic series also has potential value for crop improvement. Unlike constitutive *APO1* overexpression, regulatory editing may allow moderate increases in branch or spikelet number while retaining otherwise normal development. However, this study does not directly evaluate grain yield or possible trade-offs involving grain filling, fertility, and panicle number. The appropriate agricultural conclusion is therefore that *APO1* CNS3 provides candidate alleles for evaluating yield-related trade-offs, not that the present mutants have already been demonstrated to improve grain yield.

## Materials and methods

### Identification of CNSs in rice chromosome 6

To identify CNSs in the *APO1* regulatory region, we obtained nucleotide sequences of chromosome 6 with the *APO1* ortholog for *Leersia perrieri*, *Brachypodium distachyon*, *Aegilops tauschii*, *Hordeum vulgare*, and *Oryza sativa* Japonica from EnsemblPlants release 60 (http://plants.ensembl.org). Pairwise whole-chromosome alignments were generated between rice chromosome 6 (IRGSP-1.0) and orthologous chromosomes from the four BEP clade species using LASTZ v1.04.41 with the parameters --inner=2000 and --chain. Multiple sequence alignments of these five BEP clade species were constructed with MULTIZ/TBA v11.2; single coverage was enforced with single_cov2, and roast produced the combined alignment under the guide topology ((*O. sativa*, *L. perrieri*) (*B. distachyon*, (*A. tauschii*, *H. vulgare*))). Neutral substitution patterns were modeled by extracting fourfold degenerate (4D) sites from the rice reference annotation (IRGSP-1.0) with msa_view in PHAST v1.5 ^31^. The 4D sites were used to build a neutral phylogenetic model with phyloFit and phyloBoot, yielding a nonconserved model. For the conserved model, the chromosome 6 alignments were partitioned into 1 Mb windows using msa_split, and local conserved models were estimated with phastCons. The window-based conserved models were then averaged with phyloBoot to produce the final conserved phylogenetic model. We computed basewise conservation scores with phastCons using the neutral and conserved models learned as described above, with target coverage = 0.25 and expected conserved length = 12 bp. Conserved elements were then called as contiguous runs of positions with phastCons posterior probability ≥ 0.7 and a minimum length of 20 bp. Elements overlapping annotated coding sequences were classified as coding and excluded from downstream analyses; the remaining elements were considered CNSs.

To assess conservation at a broader phylogenetic scale, nucleotide sequences of the chromosome containing the APO1 ortholog for *Sorghum bicolor*, *Zea mays*, *Setaria italica*, and *Panicum hallii FIL2* and *Musa acuminata* were obtained from EnsemblPlants. Whole-chromosome alignments were generated for these four additional PACMAD species and *Musa acuminata* using the same LASTZ parameters, and a combined alignment of all nine Poaceae species plus Musa was constructed under the guide topology (((((*O. sativa*, *L. perrieri*) (*B. distachyon*, (*A. tauschii*, *H. vulgare*))) ((*S. bicolor*, *Z. mays*) (*S. italica*, *P. hallii*))) *Musa acuminata*)).

We surveyed the presence of CNSs in upstream and downstream of *APO1* (chr6:27480082-27481453) in *O. sativa,* and found three CNSs (CNS1: chr6:27478213-27478295; CNS2: chr6:27483827-27483993; CNS3: chr6:27484894-27485103) neighboring *APO1*. Orthologous CNSs in additional Poaceae species, including *Avena sativa OT3098, Lolium perenne, Secale cereale, Saccharum spontaneum, Echinochloa crus-galli, Digitaria exilis* and *Eragrostis tef* in EnsemblPlants, *Eremochloa ophiuroides* ^32^ and *Zoysia japonica* ^33^, *Pharus latifolius=* ^34^ and *Streptochaeta angustifolia* ^35^, as well as *Poa annua* (GCA_026929855.1), *Dactylis glomerata* (GCA_007115705.1), *Agrostis stolonifera* (GCA_052724435.1), *Holcus mollis* (PRJEB94103), *Deschampsia cespitosa* (PRJEB122840), and *Bromus sterilis* (GCA_950022295.1) from NCBI were detected by BLASTN. For *Z. mays, A. sativa, S. spontaneum, E. crus-galli, D. exilis,* and *E. tef, P. annua, A. stolonifera*, and *H. mollis*, which have undergone whole-genome duplication or polyploidization and therefore carry more than one *APO1* locus, one representative locus with the most diverged CNS2 and CNS3 was included in Figure 4b.

### Identification of orthologous CNSs and CArG box motifs

The genomic positions of CNS2 and CNS3 were examined in each species, and their location in the vicinity of the *APO1* ortholog was confirmed based on the corresponding genome annotation. CNS sequences were aligned with MAFFT v7.505. CArG boxes 1 and 2 were identified by inspection of the aligned CNS3 interval. The canonical CArG box 1 sequence (CCATTTTTG) and CArG box 2 sequence (CTGAAATG) are highly conserved across Poaceae and are consistently located in the 3’ region of CNS3. Substitutions relative to these ancestral sequences are reported in Supplementary Table 5. Cases where the motif was absent or undetectable are indicated in Supplementary Table 5. Sequence logos for CArG box 1 and CArG box 2 were generated using logomaker v1.2 ^36^ in Python. Sequences were collected from species with an identifiable motif (CArG box 1, n = 25; CArG box 2, n = 23), using one representative sequence per species. Species without an identifiable motif were excluded: *Streptochaeta angustifolia* for CArG box 1, and *S. angustifolia, Pharus latifolius*, and *Echinochloa crus-galli* for CArG box 2. For polyploid species, a single representative sequence was used. Information content at each position was calculated from the observed nucleotide frequencies relative to a uniform background.

### Vector construction for CRISPR/Cas9 genome editing

To generate genome-edited lines in the *APO1* regulatory regions, pairs of guide RNAs (gRNAs) were incorporated into each CRISPR/Cas9 vector and expressed simultaneously. Fragments containing each gRNA pair were amplified by PCR using the pUCAMdE-U3U6 vector as a template and inserted into the pZNH2GTR-U6 vector by multiple steps of enzyme digestion and ligation. The pZNH2GTR-U6 vector, derived from pZ2028 ^37^, contains a functional Cas9 cassette and spectinomycin- and hygromycin-resistance genes. Escherichia coli strain DH5α was grown in 2×YT medium containing spectinomycin at 37°C for vector cloning. Agrobacterium tumefaciens strain EHA105 was transformed with the constructed vector and grown in 2×YT medium containing spectinomycin at 28°C. The positions of the gRNA target sites are shown in Figure 2 and Supplementary Figure 2. The gRNA sequences are listed in Supplementary Table 8, and the gRNA pairs used for genome editing are listed in Supplementary Table 9.

### Identification and genotyping of edited lines

Genomic DNA was extracted from leaves of wild type and edited rice lines using the Maxwell RSC Plant DNA Kit (Promega) according to the manufacturer’s instructions. Edited *APO1* regulatory regions were amplified from genomic DNA using the primers listed in Supplementary Table 9. PCR products were separated by agarose gel electrophoresis; bands of expected size were excised and subjected to Sanger sequencing. For lines in which small indels prevented reliable band separation, genotyping was performed in the T1 generation. Homozygous alleles were confirmed by Sanger sequencing of T1 plants. Deletion or insertion junctions were determined relative to the Nipponbare reference sequence. The category, target region, and allele of each edited line are reported in Supplementary Table 1. All lines were confirmed homozygous by genotyping. Independently derived lines were obtained from independent transformation events or from T1 siblings originating from the same T0 plant.

### Rice transformation

Rice (*Oryza sativa* L. ssp. *japonica*) cv. Nipponbare served as the wild type in this study. Nipponbare seeds were de-husked, surface-sterilized with 70% ethanol and then with sodium hypochlorite 1% for 30 min, followed by 3 washes in sterile distilled water. Surface-sterilized seeds on 2N6 medium were kept at 32 °C in the dark for 4 weeks for inducing calli from seeds. The seeds were replanted in new 2N6 medium every two weeks. Agrobacterium strain EHA105 harboring the constructed vector were cultured in AB medium with spectinomycin at 23 °C in the dark for 3 days. Before the infection of the Agrobacterium, calli were cultured in new 2N6 medium at 32 °C in the dark for 3 days. Agrobacterium strain EHA105 was suspended by liquid MSL medium with acetosyringone, and added to calli. The calli were grown in N6CO medium at 23 °C in the dark. The calli were washed by sterile distilled water and meropenem trihydrate solution, and were grown in N6SE medium with meropenem trihydrate and hygromycin at 32 °C in the dark for 4 weeks. Calli were grown in ReIII medium with meropenem trihydrate and hygromycin at 28 °C under continuous light for about 6 weeks. Plants approximately 10 mm in height were grown in MSF medium at 28 °C under continuous light. Plants approximately 150 mm in height were transplanted in soil and grown in a biohazard room at 28 °C under the regime of a 14 h light/10 h dark photoperiod.

### Plant growth and panicle phenotyping

Plants used for phenotyping were grown as described above. Line 1 was a non-transgenic Nipponbare plant grown alongside the edited lines. Five plants were analyzed per line, and up to five mature panicles were collected per plant in order of emergence; panicles beyond the fifth were excluded.

For each panicle, we counted primary branches, secondary branches, total spikelets, and lateral organs. Primary branches were defined as branches arising directly from the main rachis, and secondary branches as branches arising from primary branches. Lateral-organ number was defined as the number of secondary branches plus the number of spikelets borne directly on primary branches, excluding the terminal spikelet of each primary branch. Relative secondary branching was calculated for each primary branch as the ratio of secondary branches to lateral organs, and the resulting ratios were averaged across primary branches within each panicle. Tertiary branches were recorded but were absent from all panicles examined.

For each trait, measurements from multiple panicles were averaged within each plant, and the resulting plant means were used as independent biological replicates for statistical analysis.

For the spatial analysis of lateral-organ formation, primary branches were numbered sequentially from the basal to the apical end of each panicle. Because the number of primary branches differed among panicles, the position of each primary branch was normalized as (*r −* 1)/(*N* − 1), where *r* is its basal-to-apical rank and *N* is the total number of primary branches on that panicle, as determined from the panicle summary record. The resulting coordinate ranged from 1 for the most apical primary branch to 0 for the most basal primary branch. Normalized positions were used to determine the onset position of secondary branching. For visualization and descriptive summaries, the normalized coordinate was divided into five equal-width intervals: 0–<0.2, 0.2–<0.4, 0.4–<0.6, 0.6–<0.8, and 0.8–1.0, corresponding to progressively more apical regions of the panicle. The onset position of secondary branching was defined for each panicle as the normalized position of the most basal primary branch bearing at least one secondary branch; it was treated as missing for panicles without secondary branches.

For each primary branch, we recorded the number of secondary branches, the number of spikelets borne directly on the primary branch, and the number of spikelets borne on its secondary branches. The terminal spikelet of each primary branch was included in the direct-spikelet count but was not classified as a lateral organ. Accordingly, the number of lateral organs on a primary branch was calculated as the number of secondary branches plus the number of directly borne spikelets excluding the terminal spikelet. Lateral organs were classified according to whether they developed as direct spikelets or secondary branches. Total spikelet number per primary branch was calculated as the sum of directly borne spikelets, including the terminal spikelet, and spikelets borne on secondary branches. Position-specific measurements were first calculated for each panicle and then averaged across panicles from the same plant. Plant means were treated as independent biological replicates for comparisons among lines. For summaries of mutation classes, plant means were averaged within each line, and class-level values were calculated as unweighted means of line means so that each independently derived line contributed equally. Error bars in class-level positional plots represent the standard error of the mean among line means and were not calculated for mutation classes represented by a single line.

### Statistical analysis of panicle phenotypes

Trait values were first calculated for individual panicles and then averaged within each plant. Plant means were treated as independent biological replicates in all statistical analyses; individual panicle measurements were retained for visualization but were not treated as independent replicates. When fewer than five panicles were available, the plant mean was calculated from the available measurements, and each plant contributed one value to the line-level analysis. Each line was compared with line 1 (wild type) using a two-sided Mann–Whitney U test applied separately to each of the three traits (primary branch number, total spikelet number, and secondary branches per lateral organ). P values from all pairwise tests (34 lines × 3 traits = 102 comparisons) were adjusted together using the Benjamini–Hochberg false discovery rate (FDR) procedure; adjusted p values below 0.05 were considered statistically significant. Statistical analyses were performed in Python (version 3.12) using the SciPy (version 1.13) and statsmodels (version 0.14) libraries. Exact numbers of plants and panicles contributing to each line, together with U statistics, unadjusted p values, FDR-adjusted p values, and line means, are provided in Supplementary Table 2.

For the spatial analysis, the onset position of secondary branching was compared between each line and wild type using a one-sided Mann–Whitney U test (alternative: line < wild type). Plant means of onset position of secondary branching were used as the unit of comparison. P values from all pairwise tests (34 comparisons) were adjusted together using the Benjamini–Hochberg false discovery rate (FDR) procedure. Detailed results are provided in Supplementary Table 3.

### RNA extraction and expression analysis

Total RNA was extracted from 1–2 mm young panicles from wild type and edited lines using the Maxwell RSC Plant RNA Kit (Promega) according to the manufacturer’s instructions. First-strand cDNA was synthesized from 1 μg of total RNA using the SuperScript VILO cDNA Synthesis Kit (Invitrogen) with oligo(dT) primers. qRT-PCR was performed using KOD SYBR qPCR Mix (TOYOBO) on a StepOnePlus Real-Time PCR System (Applied Biosystems). Each 20-μL reaction contained 2 μL of 20-fold-diluted cDNA, 10 μL of KOD SYBR qPCR Mix, and 0.32 μM of each primer. The thermal cycling conditions comprised an initial denaturation at 98°C for 2 min, followed by 40 cycles of 98°C for 10 s, 60°C for 10 s, and 68°C for 30 s. Amplification specificity was verified by melting-curve analysis. Primers used were as follows: APO1, 5′-TTCGCCGTCAAGAACATCT-3′ and 5′-GGGATCGAGGGAGGAGAG-3′; OsUBQ5, 5′-AACCAGCTGAGGCCCAAGA-3′ and 5′-ACGATTGATTTAACCAGTCCATGA-3′. *APO1* transcript abundance was normalized to *OsUBQ5,* and relative expression was calculated using the 2^-ΔΔCt^ method.

Associations between *APO1* transcript abundance measured by qRT-PCR and panicle traits were evaluated using genotype-level means for the eight selected genotypes. Individual plants and panicles were not treated as independent expression–phenotype pairs. Associations were tested using Spearman’s rank correlation, with Spearman’s ρ and P values reported for each trait.

### Gel shift assays

Gel shift assays were performed as previously described ^38^ with minor modifications. The coding sequences of *OsMADS14, OsMADS15, OsMADS18*, and *OsMADS34* were amplified by PCR and cloned into XhoI- and XbaI-digested pTnT vector (Promega) using the NEBuilder HiFi DNA Assembly kit (NEB). In vitro transcription and translation were carried out using 5 µg of each plasmid with the TnT SP6 High-Yield Wheat Germ Protein Expression System (Promega) in 25 µL reactions. The wild type CNS3 fragment was amplified from Nipponbare genomic DNA and cloned into the pCR-Blunt vector using the Zero Blunt TOPO cloning kit (Thermo Fisher). Mutant probes were generated by inverse PCR and self-ligation, using the plasmid containing the wild-type CNS3 fragment described above as a template. Similarly, the mut1&3 probe was generated by inverse PCR using a plasmid containing mut1 as a template and primers for mut3. Finally, using each plasmid as a template, biotin-labeled probes were synthesized by PCR using 5’-biotinylated primers annealing to the vector backbone. Primers were listed in Supplementary Table 10.

Binding reactions (20 μL total volume) were carried out in binding buffer (10 mM Tris-HCl, pH 7.5, 50 mM NaCl, 1% CHAPS, 0.5 mM EDTA, 1 mM MgCl, 0.5 mM DTT, 60 μg/mL salmon sperm DNA, and 8% glycerol), with 20 fmol of probe and 2 μL of in vitro translated protein. Samples were electrophoresed on 5% non-denaturing polyacrylamide gels pre-run at 100 V for 30 min at 4°C in 0.5× TBE buffer, followed by electrophoresis for 2 h under the same conditions. Biotin-labeled probes were detected using the LightShift Chemiluminescent EMSA Kit (Thermo Fisher).

Three sets of EMSAs were performed. First, binding of OsMADS14, OsMADS15, OsMADS18, and OsMADS34, individually and in combination, to the wild type CNS3 probe was examined. Reactions without an expression plasmid during in vitro translation were used as negative controls.

Second, the contribution of the two CArG boxes to MADS15–MADS34 binding was examined using five CNS3 probes: wild type; mut1, containing a 1-bp deletion in CArG box 1; mut2, containing a 1-bp deletion in the intervening sequence between CArG boxes 1 and 2; mut3, containing a 1-bp deletion in CArG box 2; and mut1&3, containing both the mut1 and mut3 deletions. These probes were assayed with MADS15 and MADS34.

Third, CNS3 fragments corresponding to the actual genome-edited alleles were PCR-amplified from genomic DNA using primers KT2641 and KT2642 and cloned into the pCR-Blunt vector. Biotin-labelled fragments were generated as described above, and used as EMSA probes. The assayed fragments were derived from the wild type and edited lines 28, 27, 31, 10, 30, 29, 24, 23, 26, 11, 32, and 41. Each probe was assayed in the presence or absence of MADS15 and MADS34 under the same binding conditions described above. Primer sequences used for probe preparation are listed in Supplementary Table 10.

### Validation of the *Secale cereale* CNS3 sequence

Seeds of *Secale cereale* commercially sold as ‘Raitaro’ were obtained from Takii & Co., Ltd. (Kyoto, Japan) and grown for genomic DNA extraction. Genomic DNA was extracted from leaves using the Maxwell RSC Plant DNA Kit (Promega) according to the manufacturer’s instructions. The CNS3 region containing the CArG-box was amplified by PCR using the primers listed in Supplementary Table 9 and subjected to Sanger sequencing. The obtained sequence was compared with the corresponding sequence in the Rye_Lo7_2018_v1p1p1 reference genome. The experimentally determined sequence and its discrepancy from the reference genome are shown in Supplementary Figure 4b.

### Inflorescence type classification

Inflorescence types were classified based on two structural features: spikelet attachment and branching pattern ^30^. Spikelets were categorized as sessile (directly attached), pedicellate (borne on stalks), or mixed (both). Branching was defined as either unbranched (no lateral branches) or branched (with lateral branches). Branched architectures included both hierarchically subdivided lateral axes and digitate arrangements. Three canonical inflorescence types were defined using these criteria: spike (sessile, unbranched), raceme (pedicellate, unbranched), and panicle (pedicellate, hierarchically branched). In addition, spike-like racemes and digitate inflorescences were recognized as structurally distinct forms. All classification assignments are listed in Supplementary Table 4.

### Phylogenetic comparative analysis

The species phylogeny and branch lengths were obtained from TimeTree ^39^. Inflorescence architecture was classified as branched or unbranched as described above. CNS architecture and CArG-box state were coded using the definitions in Supplementary Tables 6 and 5, respectively. The ancestral CNS configuration of core Poaceae was interpreted under parsimony: because phylogenetically distant unbranched lineages, including Pooideae wheat relatives and the PACMAD chloridoid *Zoysia japonica*, share an identical CNS2–CNS3 configuration, the conserved configuration was treated as the ancestral state for core Poaceae.

For the comparative analysis of cis-regulatory features across species, five binary features were scored for each of 24 Poaceae species and summed to generate a composite divergence score (0–5): CNS2–CNS3 spacing change (>110 bp, corresponding to the maximum CNS2–CNS3 spacing observed among unbranched species [110 bp in *Eremochloa ophiuroides*], or translocation to a different chromosome), CNS2 structural change (>30 bp), CNS3 structural change (>30 bp), CArG box 1 nucleotide substitution from the canonical motif (CCATTTTTG), and CArG box 2 change (nucleotide substitution relative to the ancestral motif CTGAAATG, deletion, or motif absence). CArG box 2 changes were further classified as minor (1–3 nucleotide substitutions relative to the ancestral motif) or major (≥4 nucleotide substitutions, deletion, or absence of an identifiable motif). Two composite divergence scores were calculated, one including both minor and major CArG box 2 changes and the other including only major changes. For polyploid species with multiple ohnologs (WGD-derived paralog), a feature was scored as changed if it was altered in at least one ohnolog.

The association between composite divergence score and inflorescence type (branched or unbranched) was tested using phylogenetic generalized least squares (PGLS) under a Brownian motion model, implemented in Python using a phylogenetic covariance matrix derived from root-to-tip and patristic distances computed from the TimeTree phylogeny. Regression coefficients were estimated by generalized least squares using SciPy (version 1.13). Individual features were tested using the same framework. Full results are provided in Supplementary Table 7.

## Data availability

Raw panicle measurements, species coding, phylogenetic trees, expression data, analysis scripts, CNS sequences across analyzed grass species, and genome-edited line sequences are available on Zenodo ^40^.

## Code availability

Analysis scripts are available on Zenodo ^40^.

## Abbreviations

CNS: Conserved noncoding sequence
APO1: ABERRANT PANICLE ORGANIZATION 1
EMSA: electrophoretic mobility shift assay
BEP: *Bambusoideae, Ehrhartoideae* and *Pooideae*
PACMAD: *Panicoideae, Aristidoideae, Chloridoideae, Micrairoideae, Arundinoideae* and *Danthonioideae*

## Acknowledgements

We thank Dr. Kuroda for providing vectors, pZNH2GTR-U6, pZNH2GTR(NG)-U6 and pUC198AMdE-U61, U62, and U63.

## Author Contributions

T.M. and J.K. conceived and designed the project. T.M., K.T., A.M., K.S., T.T., and J.K. conducted experiments. T.M. analyzed the data. T.M. wrote the first draft of the manuscript. K.T., T.T., and J.K. contributed to manuscript revision and finalization.

