## Supplementary Figures for "Recurrent evolution of an *APO1* cis-regulatory module that shapes rice panicle branching"

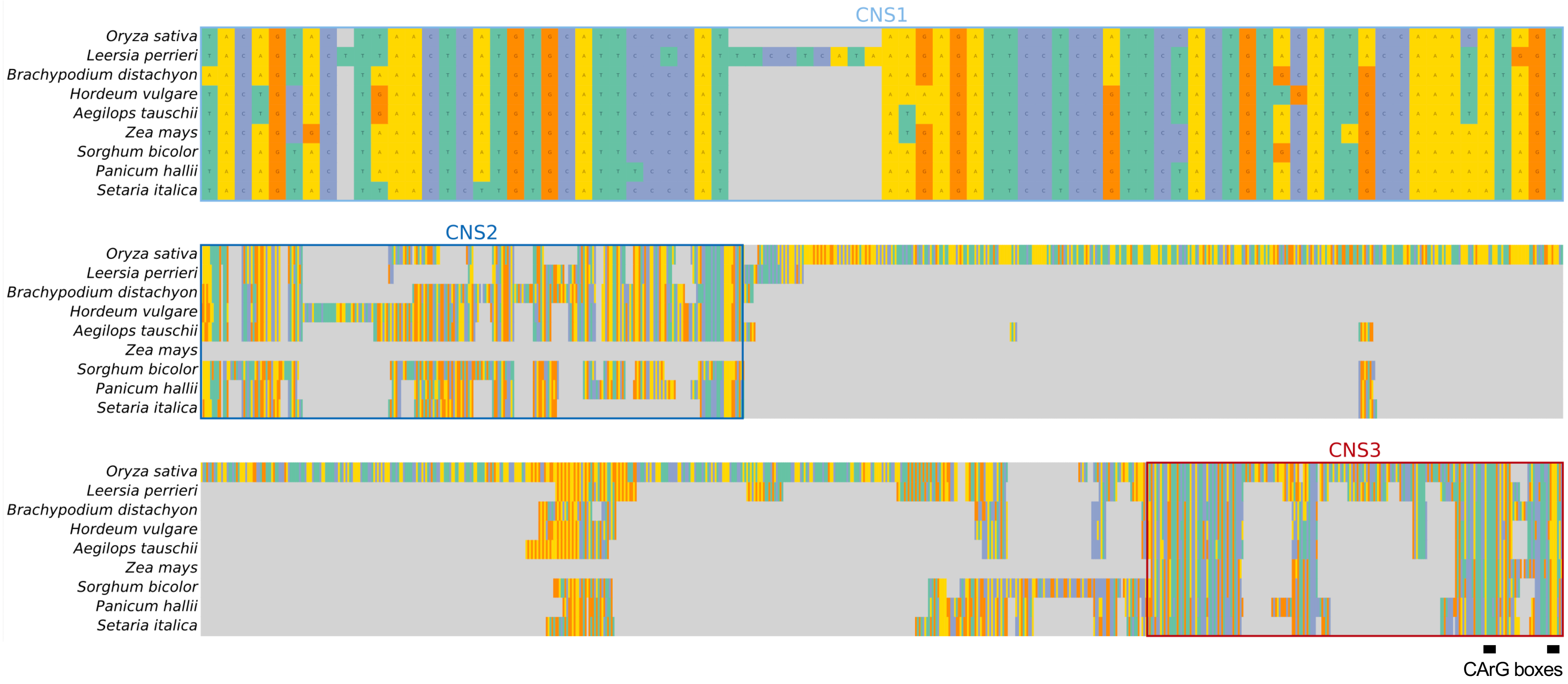


**Supplementary Fig. 1|Conservation of *APO1* cis-regulatory sequences across grasses.** Multiple-sequence alignments of CNS1, CNS2, and CNS3 in nine grass species representing the BEP and PACMAD clades. CNS1, CNS2, and CNS3 are outlined in light blue, blue, and red, respectively. Colors indicate nucleotide identity, gray denotes gaps, and black bars mark CArG boxes.

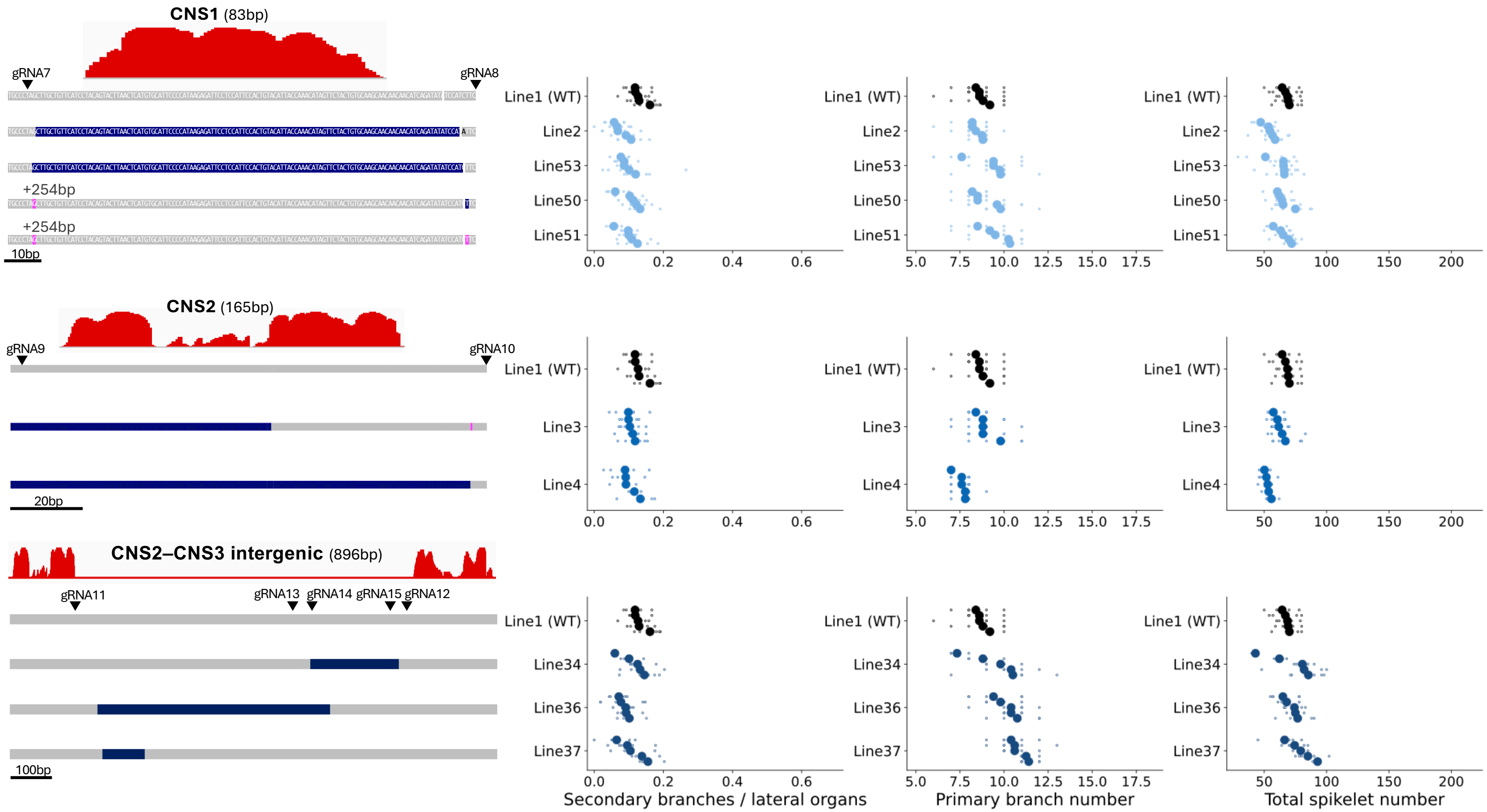


**Supplementary Fig. 2|Allele structures and panicle phenotypes of edited lines carrying mutations in the *APO1* regulatory region outside CNS3.** Edited alleles and panicle phenotypes for CNS1 (top), CNS2 (middle), and the CNS2–CNS3 intergenic region (bottom). phastCons scores, guide RNA target sites (inverted triangles), and allele structures are shown. Deletions are indicated in dark blue and insertions in pink, as in Fig. 2. Plots show secondary branches per lateral organ, primary branch number, and total spikelet number. Small open circles represent individual panicles and larger filled circles plant means (up to five panicles from each of five plants per line). Plant means were used as independent biological replicates. Plants are ordered by trait value within each line. Missing observations were not plotted or treated as zero.

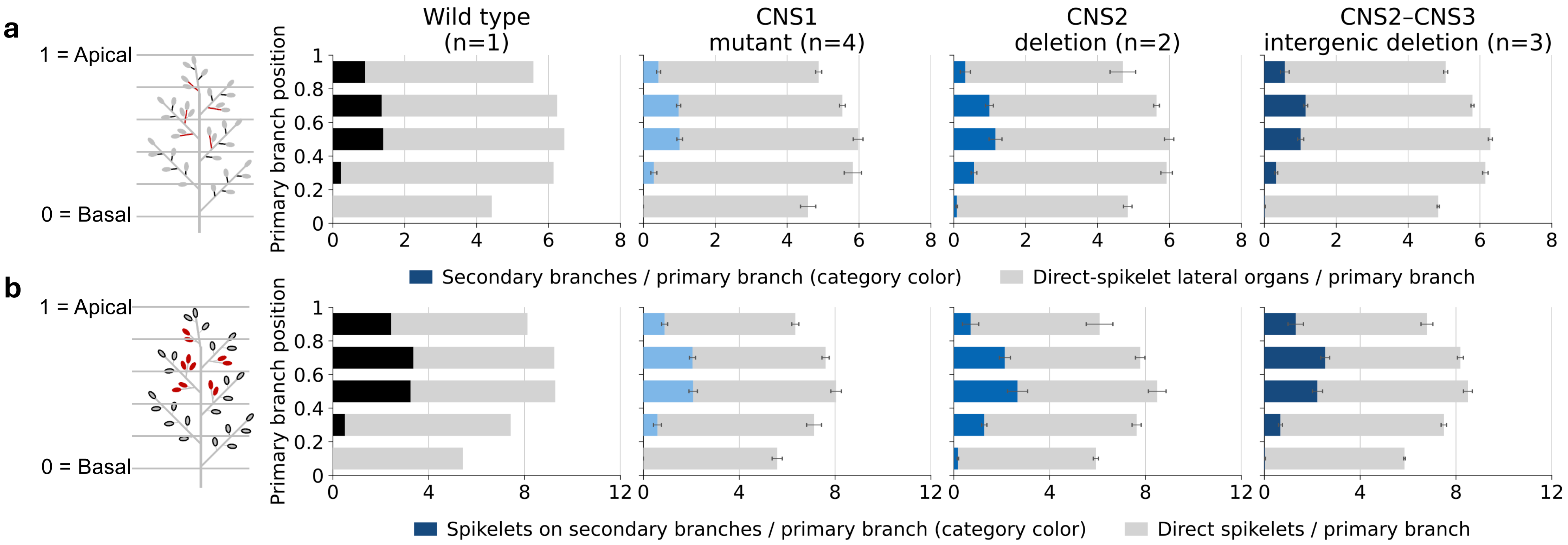


**Supplementary Fig. 3|Spatial distribution of lateral organs and spikelet production along the panicle axis in *APO1* cis-regulatory mutants. a,** Mean lateral organs per primary branch, partitioned into secondary branches (colored) and direct-spikelet lateral organs (gray), excluding terminal spikelets. **b,** Mean spikelets per primary branch, partitioned into secondary-branch-borne (colored) and direct (gray) spikelets, including terminal spikelets. **a, b,** Primary-branch positions were normalized from 0 (basal) to 1 (apical) and divided into five equal-width bins. Colors indicate mutation classes, with numbers of independent lines shown above each plot. Values are unweighted means of line means; error bars indicate SEM among line means for both components and totals.

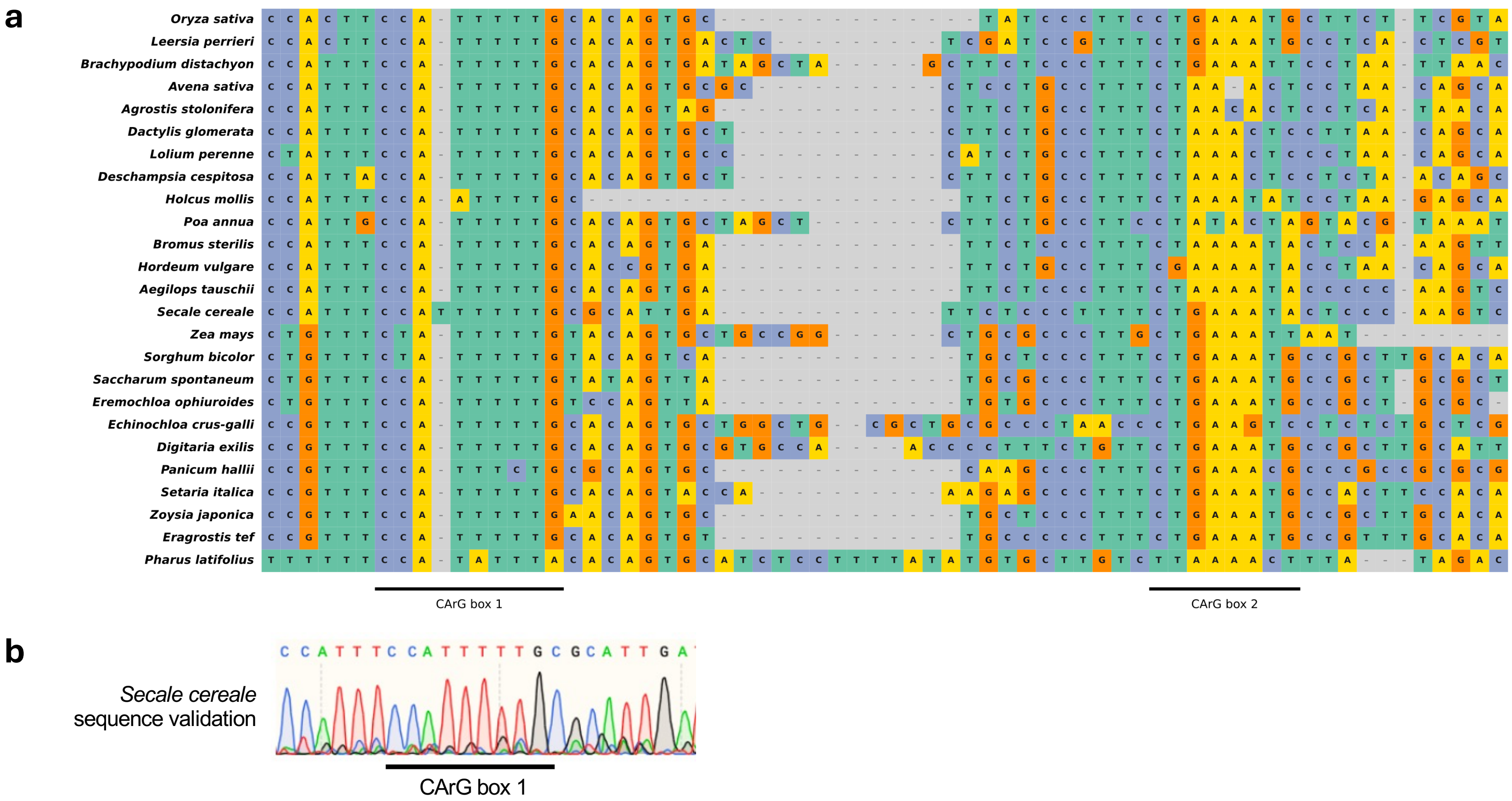


**Supplementary Fig. 4|Sequence variation in the CNS3 CArG-box region across grasses. a,** Alignment of the CNS3 region containing CArG boxes 1 and 2 across grass species with identifiable sequences, ordered as in Fig. 4b. Black bars indicate CArG boxes; colors indicate nucleotide identity and gray denotes gaps. **b,** Sanger-sequence validation of CArG box 1 in an independent Secale cereale accession. The experimentally determined sequence retained an intact motif, unlike the reference-genome sequence containing six consecutive thymine residues shown in **a**. The discrepancy may reflect an assembly error or intraspecific variation.

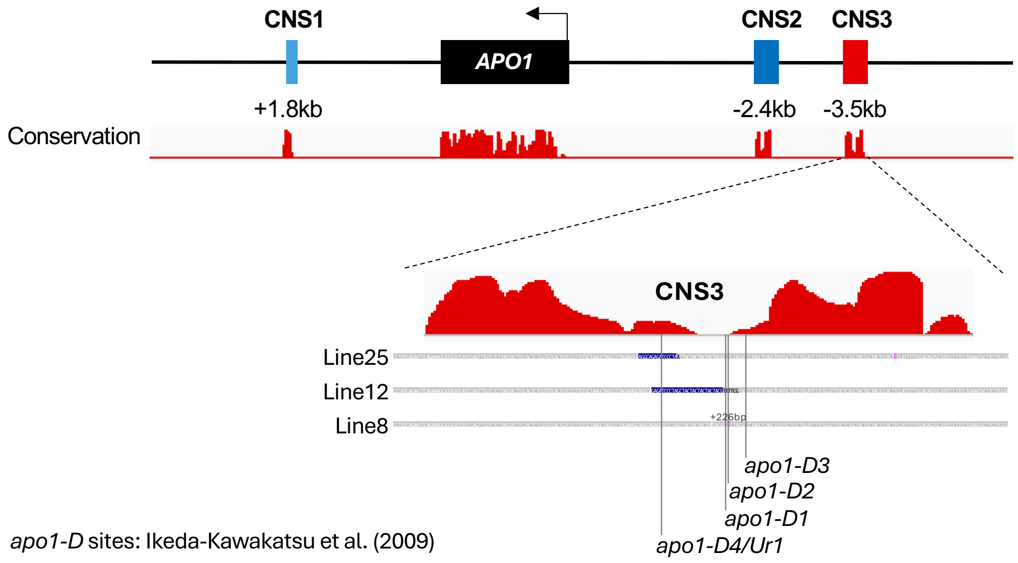


**Supplementary Fig. 5|Central CNS3 mutants and previously reported dominant *apo1-D* alleles map to a low-conservation region within CNS3.** phastCons scores across CNS3 and positions of the 226-bp insertion (line 8), deletion (line 12), and replacement (line 25), together with previously reported transposon insertion sites of *apo1-D1*, *apo1-D2*, *apo1-D3*, and *apo1-D4/Ur1*. The line 8 insertion coincides with *apo1-D2* and lies one nucleotide from *apo1-D1*; the line 12 deletion and line 25 replacement both encompass the *apo1-D4/Ur1* insertion site. All these variants map to a relatively low-conservation region of CNS3. Previously reported *apo1-D* insertion sites were redrawn from Ikeda-Kawakatsu et al. (2009).
